# Effect of zinc supplementation on certain serum biochemical parameters in growing pigs

**DOI:** 10.1101/2024.07.02.601647

**Authors:** S. Borah, S Soren, SS Deka, L Kalita, K Pame, B Borah

## Abstract

Twenty-four healthy and uniform crossbred (Hampshire × Assam Local) pigs, 4 months of age, were selected to study the effect of 500 ppm Zn supplementation on certain serum biochemical parameters for a period of four months. The animals were divided into three groups (n=8/group): group C (basal diet), group T_1_ (basal diet + 100 ppm Zn + calcium carbonate @ 1.5% DM) and group T_2_ (basal diet + 500 ppm Zn). Serum samples were collected at 15-day intervals and analysed for serum Zn, copper, total serum protein, albumin, globulin, and glucose cholesterol. The serum Zn concentration in the T_1_ group decreased significantly (P<0.01) from day 30 until day 120 of treatment, whereas in the T_2_ group, the level increased significantly (P<0.01) from day 15 onwards. The serum copper concentration did not change groups. The total protein concentration in the serum showed a significant (P<0.01) increasing trend in the T_2_ group from day 45 of treatment, and a significant (P<0.01) decreasing trend was observed in the T_1_ group compared to the C group. The serum globulin concentration decreased significantly (P< 0.05) in the T_1_ group compared with the C and T_2_ groups during the treatment period. An increasing trend in glucose and cholesterol levels was recorded at T_2_ during the treatment period. However, decrease, (P<0.01) in glucose and cholesterol levels was recorded in the T_1_ group from day 30 to day 120 of treatment. The haemoglobin concentration showed a significant (p<0.01) decreasing trend in the T_1_ group from day 45 onwards.

## INTRODUCTION

The health, productivity and reproductive performance of livestock are dependent on proper utilization of foodstuffs within the body. The utilization of different nutrients in biological systems can be studied by examining blood biochemical profiles, which also indicate health status. Among the different livestock, pigs differ from other species because of a number of biological advantages, such as high prolificacy, shorter gestation periods, shorter farrowing intervals, and better feed conversion efficiency. To achieve these advantages, pigs must meet all the nutritional requirements of their diet. Among the different nutrients, minerals play a vital role in the body due to their involvement in a large number of physiological, digestive and biosynthetic processes in biological systems. The requirements of all the essential macro- and microminerals are not widely monitored in the general management practices of pig rearing except for the requirement of iron to protect piglets from anaemia. However, other minerals needed by pigs, such as iron, calcium, and iodine, not only control a wide range of biochemical functions through their involvement in different enzyme systems in metabolic processes, e.g., Mg and Mn. Among these different minerals, Zn is an essential component of approximately 300 enzymes that are involved in a wide range of biochemical functions within the body. It also needs a continuous supply in the diet because it is not widely stored in the body. Considering the involvement of Zn and 300 enzymes in the biological system, the present study investigated the effect of supplementation with 500 ppm Zn on certain serum *biochemical parameters, viz*., total serum protein, albumin, globulin, glucose, cholesterol, etc.,, in growing pigs.

## MATERIALS AND METHODS

A total of twenty-four healthy and uniform crossbred (Hampshire × Assam Local) pigs, 4 months of age, were selected for the present experiment. The animals were divided into three groups, *viz*. The C (control), T_1_ and T_2_ treatment groups were as follows: group C was given a basal diet supplemented with 100 ppm (NRC, 1998) Zn; group T_1_ was given a basal diet supplemented with 100 ppm Zn and additional calcium carbonate @ 1.5% DM to induce Zn deficiency; and group T2 was given the same basal diet supplemented with 500 ppm Zn. The experimental animals were fed their respective diets for a period of 120 days, and the amount of feed offered was based upon the body weight of the animal according to the ICAR feeding standard for pigs (ICAR, 1990). The feed had an energy content of 2600 kcal/kg and a crude protein content of 20.29%. The animals were humanely treated, and the experiment was designed as per the guidelines of the Committee for the Purpose of Control and Supervision of Experiments on Animals (CPCSEA), Govt. of India guidelines and was approved by the Institutional Animal Ethics Committee. Ten milliliters of blood was collected from each of the experimental animals at 15-day intervals by anterior venacaval puncture, and serum samples were collected for the estimation of different biochemicals. The haemoglobin concentration was estimated immediately after collection of blood following the method of Baker et al. (1965). The serum levels of Zn and copper were estimated with an atomic absorption spectrophotometer (Varian Spectro AA220, USA) following the methods of Fick et al. (1979). Other serum biochemical measurements, such as total serum protein, albumin, globulin, glucose and cholesterol, were performed by using commercial kits manufactured by Span Diagnostics Ltd., India.

## RESULTS AND DISCUSSION

The serum Zn concentrations on day 0 of the experiment were 0.778 ± 0.03, 0.768 ± 0.03 and 0.781 ± 0.02 ppm for the C, T_1_ and T_2_ groups, respectively (Table 1). Compared with those in the T2 group, the serum Zn concentration in the T_1_ group decreased significantly (P<0.01) from 30 to 120 days of treatment, and the corresponding values for the C, T_1_ and T_2_ groups recorded on day 30 were 0.781 ± 1.02, 0.75 ± 0.01 and 0.846 ± 0.02 ppm, respectively. The opposite trend of the T_1_ group was recorded in the T_2_ group from day 15 of treatment until the end of the study. On day 120 of treatment, the serum levels of Zn recorded were 0.901 ± 0.01, 0.297 ± 0.04 and 1.07 ± 0.03 ppm for the C, T_1_ and T_2_ groups, respectively. The lowest serum Zn concentration in T_1_ might be due to the antagonistic effect of calcium on the intestinal absorption of Zn, which results in Zn deficiency (Tucker and Salmon, 1955). The induced Zn deficiency symptoms observed during the treatment period in the T_1_ group were rough hair and loss of hair over the body, most prominently in the abdominal region, nasal region and base of the tail, from day 30 onwards. Subsequently, the hairs of the abdominal region were completely lost, and the skin over the abdomen became scaly towards the end of the experiment. Some scaly stripes developed on the skin of the nasal region, and the base of the tail was completely scaly without hair. The animals were reluctant to eat, and the time taken to complete the ration was longer than that of the other groups.

**Table 1.** Serum mineral concentration (mean± S.E.) in different experimental groups during different treatment periods.

| Parameters | Treatment Group | Period of treatment (days) |  |  |  |  |  |  |  |  |
| --- | --- | --- | --- | --- | --- | --- | --- | --- | --- | --- |
|  |  | 0 | 15 | 30 | 45 | 60 | 75 | 90 | 105 | 120 |
| Serum Zn concentration (ppm) | C | 0.778<br>±<br>0.03 | 0.78<sup>a</sup><br>±<br>1.02 | 0.827<sup>a</sup><br>±<br>0.01 | 0.83<sup>a</sup><br>±<br>0.01 | 0.831<sup>a</sup><br>±<br>0.02 | 0.853<sup>a</sup><br>±<br>0.03 | 0.875<sup>a</sup><br>±<br>0.02 | 0.858<sup>a</sup><br>±<br>0.02 | 0.90<sup>a</sup><br>±<br>0.01 |
|  | T<sub>1</sub> | 0.768<br>±<br>0.03 | 0.759<sup>a</sup><br>±<br>0.01 | 0.615<sup>b</sup><br>±<br>0.03 | 0.548<sup>b</sup><br>±<br>0.03 | 0.394<sup>b</sup><br>±<br>0.05 | 0.382<sup>b</sup><br>±<br>0.02 | 0.297<sup>b</sup><br>±<br>0.02 | 0.266<sup>b</sup><br>±<br>0.02 | 0.297<sup>b</sup><br>±<br>0.04 |
|  | T<sub>2</sub> | 0.781<br>±<br>0.02 | 0.846<sup>c</sup><br>±<br>0.02 | 0.875<sup>c</sup><br>±<br>0.01 | 0.885<sup>c</sup><br>±<br>0.02 | 0.912<sup>c</sup><br>±<br>0.01 | 0.979<sup>c</sup><br>±<br>0.02 | 1.04<sup>c</sup><br>±<br>0.04 | 1.078<sup>c</sup><br>±<br>0.01 | 1.07<sup>c</sup><br>±<br>0.03 |
| Serum copper concentration (ppm) | C | 0.829<br>±<br>0.01 | 0.832<br>±<br>0.01 | 0.845<br>±<br>0.01 | 0.848<br>±<br>0.01 | 0.856<br>±<br>0.01 | 0.863<br>±<br>0.01 | 0.874<br>±<br>0.02 | 0.869<br>±<br>0.02 | 0.871<br>±<br>0.02 |
|  | T<sub>1</sub> | 0.825<br>±<br>0.02 | 0.834<br>±<br>0.01 | 0.847<br>±<br>0.01 | 0.848<br>±<br>0.01 | 0.859<br>±<br>0.01 | 0.867<br>±<br>0.04 | 0.871<br>±<br>0.03 | 0.870<br>±<br>0.02 | 0.870<br>±<br>0.02 |
|  | T<sub>2</sub> | 0.823<br>±<br>0.01 | 0.835<br>±<br>0.01 | 0.844<br>±<br>0.01 | 0.846<br>±<br>0.01 | 0.858<br>±<br>0.02 | 0.865<br>±<br>0.03 | 0.873<br>±<br>0.06 | 0.871<br>±<br>0.05 | 0.874<br>±<br>0.05 |

Means with different superscripts within a column differ significantly (P<0.01) for each parameter.

In the present study, there was no significant difference in the serum copper concentration (Table 1) between the C and T_2_ groups on different days of the treatment period. In the T_1_ group, the decrease in the bioavailability of Zn caused by the addition of calcium carbonate at a dose rate of 1.5% of DM also did not influence the serum copper concentration during different days of the treatment period. This clearly established that 100 and 500 ppm Zn in the swine diet did not have antagonistic effects on the intestinal absorption of copper. The present finding of serum copper concentrations supports the earlier reports of Buff et al. (2005) and Rincker et al. (2005) but does not support the reports of Fairweather-Tait (1995) and Lonnerdal (2000).

The total serum protein concentration (Table 2) did not significantly differ (P<0.01) between the different treatment groups until day 30 of treatment. On day 30, the total serum protein concentrations in the C, T_1_ and T_2_ groups were 6.53 ± 0.06, 6.54 ± 0.18 and 6.55 ± 0.09 g/100 ml, respectively. However, on day 45, the serum level of total protein decreased significantly (P<0.01) to a level of 6.18 ± 0.15 g/100 ml in T_1_ and increased significantly (P<0.01) in T_2_ (6.98 ± 0.07 g/100 ml) compared to that in C (6.69 ± 0.04 g/100 ml). The increasing trend in the T_2_ group and the decreasing trend in the T_1_ group continued until day 120 of treatment. On day 120 of treatment, the total serum protein concentrations in C, T_1_ and T_2_ were 6.96 ± 0.04, 6.31 ± 0.11 and 7.18 ± 0.16 g/100 ml, respectively. The elevated serum protein level in the T_2_ group following 500 ppm Zn supplementation indicated that supplemental Zn might have been involved in better assimilation of protein from available dietary protein (Grela and Pastuszak, 2004) by optimum production and activity of various proteolytic enzymes, such as carboxypeptidase A and carboxypeptidase B (Hedemann et al., 2003; Hedemann et al., 2006). This also explains the significantly (P<0.01) lower total serum protein concentration in the T_1_ group.

**Table 2.** Total serum protein and globulin (mean± S.E.) in different experimental groups during different treatment periods.

| Parameters | Treatment Group | Period of treatment (days) |  |  |  |  |  |  |  |  |
| --- | --- | --- | --- | --- | --- | --- | --- | --- | --- | --- |
|  |  | 0 | 15 | 30 | 45 | 60 | 75 | 90 | 105 | 120 |
| Total serum protein (g/100 ml) | C | 6.39<br>±<br>0.06 | 6.49<br>±<br>0.07 | 6.53<sup>a</sup><br>±<br>0.06 | 6.69<sup>a</sup><br>±<br>0.04 | 6.79<sup>a</sup><br>±<br>0.09 | 6.80<sup>a</sup><br>±<br>0.08 | 6.74<sup>a</sup><br>±<br>0.03 | 7.01<sup>a</sup><br>±<br>0.04 | 6.96<sup>a</sup><br>±<br>0.04 |
|  | T<sub>1</sub> | 6.40<br>±<br>0.05 | 6.68<br>±<br>0.06 | 6.54<sup>a</sup><br>±<br>0.18 | 6.18<sup>b</sup><br>±<br>0.15 | 6.16<sup>b</sup><br>±<br>0.13 | 6.11<sup>b</sup><br>±<br>0.06 | 6.49<sup>b</sup><br>±<br>0.16 | 6.19<sup>b</sup><br>±<br>0.08 | 6.31<sup>b</sup><br>±<br>0.11 |
|  | T<sub>2</sub> | 6.38<br>±<br>0.07 | 6.60<br>±<br>0.09 | 6.85<sup>c</sup><br>±<br>0.09 | 6.98<sup>c</sup><br>±<br>0.07 | 7.02<sup>c</sup><br>±<br>0.07 | 7.11<sup>c</sup><br>±<br>0.13 | 7.18<sup>c</sup><br>±<br>0.03 | 7.19<sup>c</sup><br>±<br>0.13 | 7.18<sup>c</sup><br>±<br>0.16 |
| Globulin (g/100 ml) | C | 3.12<br>±<br>0.03 | 3.08<br>±<br>0.04 | 3.06<sup>a</sup><br>±<br>0.05 | 3.17<sup>a</sup><br>±<br>0.04 | 3.18<sup>a</sup><br>±<br>0.01 | 3.18<sup>a</sup><br>±<br>0.03 | 3.19<sup>a</sup><br>±<br>0.07 | 3.26<sup>a</sup><br>±<br>0.09 | 3.35<sup>a</sup><br>±<br>0.09 |
|  | T<sub>1</sub> | 3.04<br>±<br>0.1 | 3.14<br>±<br>0.12 | 2.76<sup>b</sup><br>±<br>0.18 | 2.54<sup>b</sup><br>±<br>0.06 | 2.55<sup>b</sup><br>±<br>0.07 | 2.61<sup>b</sup><br>±<br>0.05 | 2.89<sup>b</sup><br>±<br>0.14 | 2.64<sup>b</sup><br>±<br>0.04 | 2.82<sup>b</sup><br>±<br>0.04 |
|  | T<sub>2</sub> | 3.07<br>±<br>0.08 | 3.30<br>±<br>0.09 | 3.25<sup>c</sup><br>±<br>0.07 | 3.26<sup>c</sup><br>±<br>0.04 | 3.23<sup>c</sup><br>±<br>0.06 | 3.28<sup>c</sup><br>±<br>0.09 | 3.32<sup>c</sup><br>±<br>0.06 | 3.49<sup>c</sup><br>±<br>0.09 | 3.48<sup>c</sup><br>±<br>0.05 |

Means with different superscripts within a column differ significantly (P<0.01) for each parameter.

In the T_1_ group, during the treatment period, the serum ALB concentration tended to increase from day 30 onwards until day 120 of the experiment. However, no such trend was observed in the control and T_2_ groups during the study period. When comparing the values of serum ALB among the different treatment groups, no significant differences were recorded. However, markedly greater values of serum ALB were recorded in the T_1_ group than in the C and T_2_ groups. The present study revealed that neither Zn supplementation nor induced Zn deficiency affected the serum ALB concentration.

The serum globulin concentration (Table 2) in the T_1_ group showed a significant decreasing trend from 30 days (2.76 ±0.18 g/100 m) to 120 days (2.82±0.04 g/100 ml), possibly due to improper absorption and metabolism of dietary protein (Hedemann et al., 2003; Hedemann et al., 2006). However, the T_2_ group showed a significant (P<0.01) increasing trend from day 30 onwards until the end of the experiment, indicating that supplementation with Zn has a stimulatory effect on the development of the immune system (Rupic et al., 1998; Azizzadeh et al., 2005) in pigs. The serum Gamma globulin concentrations on days 0, 15, 30 and 120 of the treatment periods in the C group were 0.67 ± 0.03, 0.72 ± 0.03, 0.76 ± 0.03, and 1.06 ± 0.04 g/100 ml, respectively. The corresponding values recorded for the T_1_ group were 0.68 ± 0.02, 0.74 ± 0.03, 0.70 ± 0.02, and 0.84 ± 0.02 g/100 ml, and those for the T_2_ group were 0.67 ± 0.02, 0.90 ± 0.05, 0.95 ± 0.03, and 1.31 ± 0.05 g/100 ml, respectively. The serum Gamma globulin concentration decreased significantly (P< 0.01) from day 30 to day 120 of treatment in the T_1_ group, while a significant (P< 0.01) increasing trend was recorded in the C and T_2_ groups from day 15 to day 120 of treatment. Higher values of gamma globulin were recorded for the C and T_2_ groups, indicating that Zn supplementation has a stimulatory effect on immune development through maintenance of the function of lymphoid organs, T-helper cells, and thymic hormones to stimulate the activity of the thymus gland (Golden et al., 1977; Fernandes et al., 1979). A decreased level of An increasing trend in blood glucose levels (Table 3) was observed at T_2_ during the treatment period, but the increasing trend was not statistically significant compared to that in the C group. However, a significant (P<0.01) decrease in the blood glucose level was detected in the T_1_ group beginning on day 30 (78.56±0.78 mg/100 ml), which decreased to 76.01±1.28 mg/10 ml on day 120 of treatment. The lowest level of blood glucose might be due to the lower activity of carbohydrate-digesting enzymes, which are dependent on the dietary level of Zn in pigs (Hedemann et al., 2006), and to lower food intake due to the induced deficiency of Zn, as observed in the experiment. A higher level of blood glucose in the T_2_ group indicates better utilization of dietary carbohydrates by digestive enzymes and is the cause of the higher concentration of blood glucose in this group.

**Table 3.** Serum glucose and cholesterol levels (mean± S.E.) in different experimental groups during different periods of treatment.

| Parameters | Treatment Group | Period of treatment (days) |  |  |  |  |  |  |  |  |
| --- | --- | --- | --- | --- | --- | --- | --- | --- | --- | --- |
|  |  | 0 | 15 | 30 | 45 | 60 | 75 | 90 | 105 | 120 |
| Glucose (mg/100 ml) | C | 79.39<br>±<br>1.87 | 79.88<br>±<br>1.25 | 79.47<sup>a</sup><br>±<br>1.08 | 79.35<sup>a</sup><br>±<br>1.69 | 80.60<sup>a</sup><br>±<br>1.21 | 80.41<sup>a</sup><br>±<br>0.75 | 79.98<sup>a</sup><br>±<br>0.98 | 80.84<sup>a</sup><br>±<br>1.33 | 79.89<sup>a</sup><br>±<br>1.13 |
|  | T<sub>1</sub> | 79.18<br>±<br>1.18 | 79.35<br>±<br>0.79 | 78.56<sup>b</sup><br>±<br>0.78 | 77.14<sup>b</sup><br>±<br>0.84 | 77.02<sup>b</sup><br>±<br>1.09 | 77.00<sup>b</sup><br>±<br>0.84 | 76.48<sup>b</sup><br>±<br>1.05 | 76.47<sup>b</sup><br>±<br>1.19 | 76.01<sup>b</sup><br>±<br>1.28 |
|  | T<sub>2</sub> | 80.14<br>±<br>0.99 | 80.82<br>±<br>0.44 | 80.14<sup>a</sup><br>±<br>1.23 | 81.67<sup>a</sup><br>±<br>0.91 | 82.43<sup>a</sup><br>±<br>0.63 | 82.09<sup>a</sup><br>±<br>0.90 | 82.58<sup>a</sup><br>±<br>1.09 | 83.21<sup>a</sup><br>±<br>1.72 | 82.86<sup>a</sup><br>±<br>1.57 |
| Cholesterol (mg/100 ml) | C | 169.36<br>±<br>5.09 | 171.36<br>±<br>5.64 | 175.28<br>±<br>4.99 | 177.26<sup>a</sup><br>±<br>6.96 | 177.01<sup>a</sup><br>±<br>6.77 | 177.98<sup>a</sup><br>±<br>5.38 | 179.89<sup>a</sup><br>±<br>4.86 | 180.45<sup>a</sup><br>±<br>3.89 | 180.86<sup>a</sup><br>±<br>9.55 |
|  | T<sub>1</sub> | 169.84<br>±<br>6.03 | 169.98<br>±<br>3.59 | 168.89<br>±<br>8.14 | 169.25<sup>b</sup><br>±<br>6.92 | 171.03<sup>b</sup><br>±<br>2.44 | 169.47<sup>b</sup><br>±<br>9.04 | 163.09<sup>b</sup><br>±<br>3.88 | 164.12<sup>b</sup><br>±<br>6.70 | 162.29<sup>b</sup><br>±<br>9.48 |
|  | T<sub>2</sub> | 170.17<br>±<br>6.78 | 172.22<br>±<br>8.37 | 174.47<br>±<br>2.07 | 177.51<sup>a</sup><br>±<br>7.95 | 179.29<sup>a</sup><br>±<br>2.96 | 179.73<sup>a</sup><br>±<br>5.84 | 180.32<sup>a</sup><br>±<br>4.23 | 182.41<sup>a</sup><br>±<br>7.16 | 182.53<sup>a</sup><br>±<br>4.99 |

Means with different superscripts within a column differ significantly (P<0.01) for each parameter.

The serum level of cholesterol (Table 3) was not affected by supplementation with 500 ppm Zn in the T_2_ group compared to the C group. However, the T_1_ group had significantly (P<0.01) lower (169.25±6.92 mg/100 ml) serum cholesterol concentrations on day 30 of treatment, and the cholesterol concentration further decreased to 162.29±9.48 mg/100 ml on day 120 of treatment. This might be due to the lower availability of blood glucose in the T_1_ group, as evidenced by the present study because glucose serves as a source for the synthesis of cholesterol in the biological system.

A steady increase in the level of haemoglobin was recorded in the C and T_2_ groups during the experimental periods. However, a significant (p<0.01) decreasing trend was noted in the T_1_ group from day 45 (10.90±0.18 g/100 ml) to day 120 (9.10±0.18 g/100 ml). Her haemoglobin level on day 45 in C and T_2_ was 11.15 ±0. 15 and 12.21±0.05 g/100 ml, respectively. The haemoglobin concentrations on day 120 of treatment were 11.28±0.19 and 12.68 ±0.45 g/100 ml for the C and T_2_ groups, respectively. The decreased concentration of haemoglobin in the T_1_ group might be because the enzyme aminolaevulinate dehydratase required for bodily synthesis of haem is Zn dependent (Warren et al., 1998), and Zn is also associated with the synthesis of **erythrocytes** (Nishiyama et al., 1998; Rupic et al., 1998; Hassan et al., 2001) and might be due to improper metabolism of dietary protein for synthesis of the globin part of haemoglobin.

## Notes

### Competing Interest Statement

The authors have declared no competing interest.

## REFERENCES

Azizzadeh, M.; Mohri, M. and Seifi, H.A. 2005. Effect of oral Zn supplementation on hematology, serum biochemistry, performance, and health in neonatal dairy calves. Comparative Clinical Pathology, 14 :67–71.

Baker, E.J.; Silverton, R.E. and Luckcock, E.D. 1965. An Introduction to Medical Laboratory Technology 1st Edn. Butterworth and Co. (Publisher) Ltd. London.

Buff, C.E.; Bollinger, D.W.; Ellersieck, M.R.; Brommelsiek, W.A. and Veum, T.L. 2005. Comparison of growth performance and Zn absorption, retention, and excretion in weanling pigs fed diets supplemented with Zn-polysaccharide or Zn oxide. J. Anim. Sci. 83:2380–2386.

Fairweather-Tait, S.J. 1995. Iron-Zn and calcium-Fe interactions in relation to Zn and Fe absorption. Proc. Nutr. Soc. 54:465–473.

Fernandes, G.A.; Nair, K.; Onoe, R.; Tanaka, R.F. and Good, R.A. (1979). Impairment of Cell-mediated Immunity Functions by Dietary Zn Deficiency in Mice. Proc. Natl. Acad. Sci. USA, 76: 457.

Fick, K.R.; McDowel, L.R.; Miles, P.H.; Wilkilson, N.S.; Funk, J.D.R and Conrad, J.H. 1979. Methods of Mineral Analysis for Plant and Animal Tissues. 2nd edn., Deptt. Anim. Sci. University of Florida, Gainesville.

Grela, E.R and Pasuszak, J. (2004). Nutritional and prophylactic importance of Zn in pig production. Medycyna weterynaryjna., 60: 1254–1258.

Golden, M.H.N.; Jackson, A.A. and Golden, B.E. (1977). Effect of Zn on Thymus of Recently, Mal-nourished Children. Lancet, 2:1, 057.

Hassan, A. El Hendy; Mokhtar, I. Yousef and Nasser, I. Abo El-Naga (2001). Effect of dietary Zn deficiency on hematological and biochemical parameters and concentrations of Zn, copper, and iron in growing rats. Toxicology, 167 : 163–170.

Hedemann, M.S.; Hojsgaard, S. and Jensen, B.B. 2003. Small intestinal morphology and activity of intestinal peptidases in piglets around weaning. J. Anim. Physiol. Anim. Nutr., 87:32–41.

Hedemann, M.S.; Jensen, B.B. and Poulsen, H.D. (2006). Influence of dietary Zn and copper on digestive enzyme activity and intestinal morphology in weaned pigs. J. Aniam. Sci., 84: 3310–3320.

I.C.A.R. 1990. Handbook of animal husbandry. ICAR, New Delhi, pp-146-158.

Lonnerdal, B. 2000. Dietary factor influencing Zn absorption. J.Nutr.130:1378–1383.

National Research Council. 1998. Nutrient Requirements of Swine. 10th Ed. National Academy Press, Washington, DC.

Nishiyama, S.; Kozo, I.; Tadashi, M.; Akimasa, H. and Ichiro, M. 1998. Zn Status Relates to Hematological Deficits in Middle-Aged Women. Journal of the American College of Nutrition, 17: 291–295.

Pekarek, R.S.; Sandstead, H.H.; Jacob, R.A. and Barcome, D.F. 1979. Abnormal cellular immune response during acquired Zn deficiency. Am. J.Clin. Nutr., 32: 1466–1471.

Rupic, V.; Libuka, I.L; Svjetlana, C.M. and. Boac, R. 1998. Plasma proteins and haematological parameters in fattening pigs fed different sources of dietary Zn. Acta. Veterinaria Hungerica, 46: 111–126.

Rincker, M.J.; Hill, G.M.; Link, J.E.; Meyer. A.A. and Rowntree, J.E. 2005. Effect of dietary Zn and iron supplementation on mineral excretion, body composition and mineral status of nursery pigs. J. Anim. Sci. 83:2762–2774.

Snedecor, G.W. and Cochran, W.G. 1994. Statistical Methods, 8th edn. The IOWA State University Press, IOWA.

Tucker, H.F and Salmon, W.D. 1955. Parakeratosis of Zn deficiency disease in the pig. Proc. Soc. Exp. Biol. Med., 88: 613.

Warren, M.J.; Cooper, J.B.; Wood, S.P. and Peter, M.S. 1998. Lead poisoning, haem synthesis and 5-aminolaevulinic acid dehydratase. Trends in Biochemical Sciences, 23 : 217–221.

